# Transcriptional profiling reveals running promotes cerebrovascular remodeling in young but not aged mice

**DOI:** 10.1101/574673

**Authors:** Kate E. Foley, Stanley Yang, Leah C. Graham, Gareth R. Howell

## Abstract

**Background:** The incidence of dementia and cognitive decline is increasing with no therapy or cure. One of the reasons treatment remains elusive is because there are various pathologies that contribute to age-related cognitive decline. Specifically, with Alzheimer’s disease, targeting to reduce amyloid beta plaques and phosphorylated tau aggregates in clinical trials has not yielded results to slow symptomology, suggesting a new approach is needed. Interestingly, exercise has been proposed as a potential therapeutic intervention to improve brain health and reduce the risk for dementia, however the benefits throughout aging are not well understood.

**Results:** To better understand the effects of exercise, we preformed transcriptional profiling on young (1-2 months) and midlife (12 months) C57BL/6J (B6) male mice after 12 weeks of voluntary running. Data was compared to age-matched sedentary controls. Interestingly, the midlife running group naturally broke into two cohorts based on distance ran - either running a lot and more intensely (high runners) or running less and less intensely (low runners). Midlife high runners had lower LDL cholesterol as well as lower adiposity (%fat) compared to sedentary, than midlife low runners compared to sedentary suggesting more intense running lowered systemic markers of risk for age-related diseases including dementias. Differential gene analysis of transcriptional profiles generated from the cortex and hippocampus showed thousands of differentially expressed (DE) genes when comparing young runners to sedentary controls. However, only a few hundred genes were DE comparing either midlife high runners or midlife low runners to midlife sedentary controls. This indicates that, in our study, the effects of running are reduced through aging. Gene set enrichment analyses identified enrichment of genes involved in extracellular matrix (ECM), vascular remodeling and angiogenesis in young runners but not midlife runners. These genes are known to be expressed in multiple vascular-related cell types including astrocytes, endothelial cells, pericytes and smooth muscle cells.

**Conclusions:** Taken together these results suggest running may best serve as a preventative measure to reduce risk for cerebrovascular decline. Ultimately, this work shows that exercise may be more effective to prevent dementia if introduced at younger ages.

## Introduction

With an aging population, the impact of age-related cognitive impairment is increasing (1) (2) (3) (4). Cognitive impairment can involve many symptoms, including memory loss, newfound difficulties with language, and inability to make every-day decisions (5). Further, the incidence of many dementias for which aging is the major risk factor, such as Alzheimer’s Disease (AD), affecting approximately 5.4 million people in the year 2016, and expected to increase to over 13 million in the next few years (6). For example, the majority of clinical trials for AD therapeutics have focused on reducing hallmark pathologies such as amyloid beta accumulation and tau tangle aggregates. However, these pathologies do not always correlate to cognitive function, and therefore these therapeutic targets may not rescue the initial symptoms that burden the patient (7) (8). Currently, despite numerous preclinical studies and clinical trials, there are no therapies for cognitive decline.

Non-pharmacological interventions have been proposed as alternatives to pharmacological treatments to prolong brain health – reducing risk for age-related cognitive decline and dementias. Obesity and physical inactivity increase risk for cognitive decline and dementia, suggesting that interventions such as diet and exercise can mitigate risk (9). Exercise has positive effects, not just improving systemic health, but also cerebral health through increases in cerebral plasticity, neurogenesis, as well as hippocampal and cortical volume (10, 11). For example, exercise earlier in life correlated with reduced cognitive impairment with age (12). The cerebral benefits of exercise may arise through the induction of brain derived neurotropic factor (BDNF), which triggers neuronal proliferation in the dentate gyrus (13). A recent study has also explored the use of running, as well as an AAV-based gene therapy to increase neuronal proliferation and survival, in a mouse model of AD (14). Viral-induced neurogenesis alone did not benefit cognition as well as running did, suggesting that running also promotes non-neuronal changes that improve cognition. Therefore, the full spectrum of processes by which running promotes brain health and reduces risk for dementias remains unclear. Furthermore, given that many such studies are performed in young mice, it remains unknown whether the beneficial effects of running persist through multiple life stages.

Here, we chose an unbiased transcriptional profiling approach to better understand the effects of running on overall brain health. To date, an extensive evaluation of the transcriptome of the brain in response to running across ages has not been assessed. RNA sequencing was performed on the cortex and hippocampus from young and middle aged (midlife) C57BL/6J (B6) male mice that were provided running wheels for 12 weeks. Transcriptional profiles were compared to aged-matched sedentary controls. Within the midlife running cohort, half the mice ran markedly faster and farther (high runners) than the other half (low runners). This provided natural variation in our midlife running dataset and allowed us to also interrogate how the intensity of running impacted systemic bodily health as well as transcriptional profiles in the brain. The young cohort did not exhibit such variation in their voluntary exercise. Transcriptomes of young running mice showed considerably more differentially expressed (DE) genes compared to either low-running or high-running midlife cohorts. Gene set enrichment analyses revealed enrichment of genes in pathways implicated in vascular remodeling in young, but not midlife mice.

## Materials and Methods

### Mouse Strains

All experiments involving mice were conducted with approval and accordance described in the Guide for the Care and Use of Laboratory Animals of the National Institutes of Health. All experiments were approved by the Animal Care and Use Committee at The Jackson Laboratory. All mice used in this study were male C57BL/6J (B6, stock number JR00664, The Jackson Laboratory). Only males were used for these experiments due to the confounding variability of the estrus cycle in female mice. Further experimentation is needed to evaluate how the estrus cycle may influence the effects of running and running with age. Mice were kept in a 12/12-hour light/dark cycle and fed ad libitum 6% kcal fat standard mouse chow.

### Exercise by Voluntary Running

Group housed mice (two-three per pen) were provided access to low profile saucer wheels (Innovive Inc) 24 hours a day for 12 weeks. Sedentary mice did not have access to running wheels. In the first experiment, young mice were housed from wean (1-2mo) until the end of the experiment (4-5mo). In the second experiment, mice were aged without wheel access, and at 12mo were provided running wheels. For both experiments, in the final week, mice were individually housed and given a trackable low-profile running wheel (Med Associates Inc.). Running wheel rotations were measured in 1-minute bins to allow for distance traveled (sum of rotations) calculated per mouse each night. Percent of time at each speed was calculated by totaling the number of minute bins that mice ran between 0, 1-30 rotations, 31-70 rotations, 71-100 rotations and 100+ rotations and dividing by the total amount of minutes tracked.

### Nuclear Magnetic Resonance (NMR) Imaging

The midlife running cohort was subjected to NMR imaging one week before harvest. Weight was taken and mice were briefly placed into a Plexiglas tube 2.5 inches by 8 inches which was then subjected to NMR (EchoMRI, Houston, TX). Magnetic field was measured by a 5-gauss magnet. Measurements included weight, lean muscle mass, fat mass, and water composition. Adiposity was calculated by (fat/body weight) × 100. Percentage lean muscle mass was calculated by (lean muscle mass/body weight) × 100.

### Harvesting, Tissue Preparation and Blood Chemistry

All mice were euthanized by intraperitoneal injection of a lethal dose of Ketamine (100mg/ml)/Xylazine(20mg/ml) and blood was collected at harvest through approved cardiac puncture protocols. Mice were perfused intracardially with 1X PBS. Brains were carefully dissected and hemisected sagittally. Hippocampus (Hippo) and cortex (Ctx), were then carefully separated and snap frozen in solid CO_2_ for RNA-sequencing. Blood was also collected in K2 EDTA (1.0mg) microtainer tubes (BD) at harvest (non-fasted) and kept at room temperature for at least 30 minutes to prevent clotting and then centrifuged at 22°C for 15 minutes at 5000rpm. Serum was carefully collected and aliquoted. Serum was characterized on the Beckman Coulter AU680 chemistry analyzer.

### RNA and Protein Extraction, Library Construction and RNA Sequencing

RNA sequencing (RNA-seq) was performed by The Jackson Laboratory Genome Technologies Core. RNA extraction involved homogenization with TRIzol (Invitrogen) as previously described (15). RNA was isolated and purified using the QIAGEN miRNeasy mini extraction kit (QIAGEN) in accordance with manufacturer’s instructions. RNA quality was measured via the Bioanalyzer 2100 (Agilent Technologies) and poly(A) RNA-seq sequencing libraries were compiled by TruSeq RNA Sample preparation kit v2 (Illumina). Quantification was performed using qPCR (Kapa Biosystems). RNA-seq was performed on the HiSeq 4,000 platform (Illumina) for 2×100bp reads for a total of 45 million reads according to the manufacturer’s instructions.

### RNA-seq quality control and gene set enrichment

Quality control for each sample was completed using NGSQCToolkit v2.3 which removed adaptors and trimmed low quality bases (Phred<30) (16). To quantify gene expression of the trimmed reads, we used RSEM v1.2.12 which uses Bowtie2 v2.2.0 for alignment of these reads (17). We used mouse genome mm-10 based upon the B6 reference genome. Differential gene expression (DGE) analysis was completed on the hippocampus and cortex separately, using EdgeR 3.20.9 (18). A second quality control step included filtering out genes with less than at least 1 read per million for more than one sample. Normalization of trimmed mean of M values (TMM) was performed and quasi-likelihood F-test was used to attain DGE. Differentially Expressed genes (DE genes) were identified by comparing (i) young running to young sedentary, (ii) midlife low running to midlife sedentary, and (iii) midlife high running to midlife sedentary for each tissue (cortex or hippocampus). Genes were considered DE if the False Discovery Rate was less than 0.05 (FDR<0.05). Given the young and midlife running were run at separate times it was not possible to directly compare the young data to the midlife data.

Ingenuity Pathway Analysis (IPA) was used to identify enriched canonical pathways for each DE gene list. Additionally, Database for Annotation, Visualization and Integrated Discovery (DAVID, v6.8) was used to identify enrichment of Kyoto Encyclopedia of Genes and Genomes (KEGG) pathways and Gene Ontology (GO) terms, with the background genesets being all trimmed normalized genes for each comparison. Enriched KEGG pathways and GO terms with FDR <0.05 were considered significant. Cancer-related pathways were excluded from visualization.

### Statistical Analyses

Details of statistical analyses of RNA-seq data is provided above. All other statistical analyses were performed in GraphPad Prism v7.0a. Body composition and lipid profiling results between the young running and young sedentary cohorts utilized an unpaired t-test. Body composition and lipid profiling results between midlife sedentary, low runners, and high runners were compared with a one-way ANOVA.

## Results

### Voluntary running distances at midlife showed a bimodal distribution

To understand the molecular changes in the brain in response to voluntary running, running wheels were provided to young (1-2 month old, mo) and midlife (12 mo) B6 male mice for 12 weeks. Age-matched sedentary controls had no access to a running wheel **(Fig. 1A)**. To quantify running, wheel rotations per minute were assessed overnight, when mice are most active, for at least five nights during the last week of the experiment. Although there was some variation between average wheel rotations per night in the young cohort, five of six mice averaged over 10,000 rotations per night. Of the 12 midlife running mice however, half averaged fewer than 10,000 rotations per night, deemed ‘Low Runners’, and half averaged greater than 10,000 rotations per night, deemed ‘High Runners’ (**Fig. 1B)**. This was not due to dominance in group housing (**Table S1)**. High runners ran a similar distance (on average, greater than 10,000 rotations per night) to young runners. Additionally, midlife high runners spent more time running compared to midlife low runners (**Fig. 1C)**. High runners also were quantified to have run at 100 rotations per minute, while the low runners did not show this ability. Young sedentary mice gained more weight over the course of the experiment than young running mice (**Fig. 1E)**. Similarly, in the midlife running cohort, high runners tended to weigh less throughout the experiment, although not significantly at the conclusion of the 12 weeks (**Fig. 1F)**. To better understand body composition in the midlife cohort, mice were subjected to NMR a week before harvest to assess fat and lean mass percentages. Midlife high runners showed a significant decrease in fat mass (g) reflected by a corresponding decrease in adiposity (%fat) (**Fig. S1A, S1B, S1D).** Lean muscle mass was not different among the groups but the midlife high running cohort had a higher percent lean muscle mass compared to low running and sedentary cohorts, presumably due to the lower body weight of the high runners (**Fig. S1C, S1E)**. This indicates that weight loss of the high run cohort was primarily fat mass.

**Figure 1:**
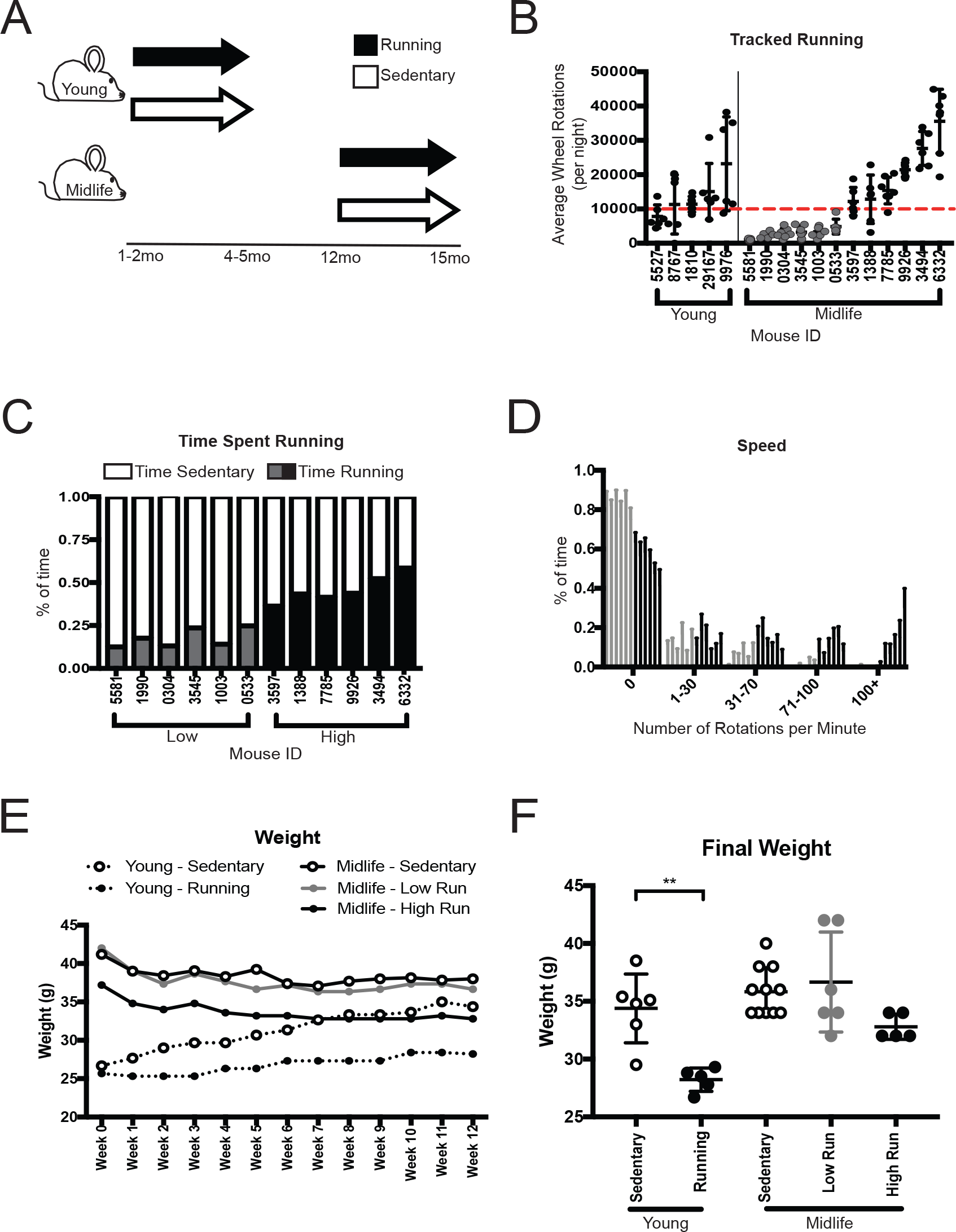
Voluntary Running at young and midlife reveals natural variation in running intensity. (A) Experimental strategy of running mice. Young mice were given access to voluntary running wheels for 12 weeks starting at 1-2 months of age and ending at 4-5 months of age. Midlife mice were given access to wheels for 12 weeks starting at 12 months of age and ending at 15 months of age. (B) Average wheel rotations per night showed dichotomous response of midlife running. Midlife mice that ran below 10,000 rotations per night (red line), were deemed ‘Low’ (grey). Midlife mice that ran above 10,000 rotations per night, deemed ‘High’ (black). (C) Low runners spent less time (% of time) running than high runners. (D) Midlife High runners ran faster, with a higher percentage of time running above 100 rotations per minute. (E) Weight over the course of running experiment. (F) Final weight at week 12 showed a significant difference in young sedentary and young runners (**p =0.0018).

To assess whether running during midlife altered metabolic indicators of health in the blood, we measured cholesterol composition, non-fasted glucose, triglycerides, and non-essential fatty acids (NEFA) levels. Total cholesterol was significantly reduced in young runners compared to age-matched sedentary controls (**Fig. 2A)**. Midlife high runners also showed a significantly lower total cholesterol profile compared to midlife sedentary controls, which could be attributed to the decrease in LDL cholesterol (**Fig. 2A-2C)**. HDL cholesterol remained unchanged with running **(Fig. 2C)**. Non-fasted glucose levels were not significantly different, although there was trend towards lower blood glucose in running compared to sedentary mice at both ages (**Fig. 2D)**. Triglycerides and non-essential fatty acids (NEFA) were reduced by running at a young age, but this effect was not seen in the midlife cohort (**Fig. 2E, 2F)**. Taken together, there was a significant shift towards a healthier body composition in midlife high runners compared to low runners. Further, systemic health benefits such as levels of total cholesterol, including LDL, can be altered at midlife due to more intense running.

**Figure 2:**
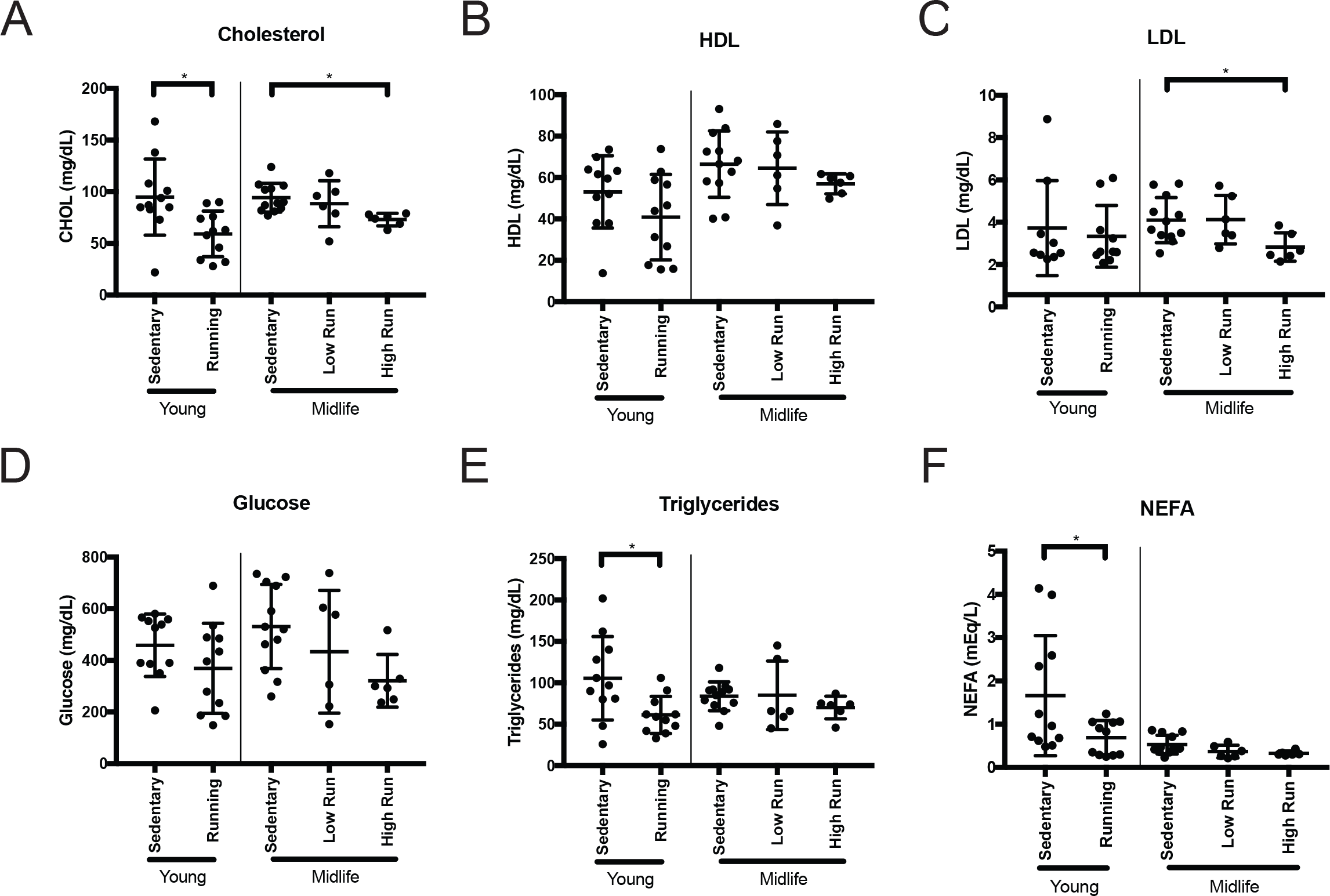
Lipid profiling of the blood showed ability to alter cholesterol composition in midlife. (A) Significant difference in total cholesterol serum concentration at harvest in young runners and between midlife high runners to midlife sedentary mice (young *p= 0.0124, midlife *p= 0.0282). (B) No significant difference in High Density Lipoprotein (HDL) serum concentration between runners and sedentary mice. (C) Significant difference in Low Density Lipoprotein (LDL) serum concentration between midlife high runners and midlife sedentary mice (*p= 0.0489). (D) No change in non-fasted glucose serum concentration across all cohorts. (E) Significant decrease in in serum triglyceride between young run and young sedentary concentration (*p= 0.0151). (F) Significant reduction in Non-Essential Fatty Acid (NEFA) serum concentrations between young run and young sedentary cohorts (*p= 0.0374).

### Young running affects transcriptional signatures more robustly than midlife running

To assess transcriptional changes in the brain after running at two different ages, RNA-seq was performed on hippocampus and cortex – vulnerable regions in age-related cognitive decline and dementia **(Fig. S2, S3)**. Differentially expressed (DE) genes (FDR <0.05) were identified by comparing (i) young running to young sedentary, (ii) midlife high runners to midlife sedentary, and (iii) midlife low runners compared to midlife sedentary (**Table S2-8,** see methods). In the cortex, there were 1,252 DE genes (742 upregulated, 510 downregulated) when comparing young runners to young sedentary mice (**Fig. 3A, 3B)**. However, there were only 214 DE genes (70 upregulated, 144 downregulated) in the midlife high runners compared to midlife sedentary, and 67 DE genes (20 upregulated and 47 downregulated) comparing midlife low runners to midlife sedentary (**Fig. 3B)**. Hippocampal analysis revealed similar results to the cortex. In young mice there were 2,026 DE genes (1,498 upregulated, 528 downregulated) in the young runners compared to young sedentary (**Fig. 3F, 3G)**. In midlife mice, there were 271 DE genes (87 upregulated, 184 downregulated) comparing midlife high runners to midlife sedentary controls, and 172 DE genes (80 upregulated, 92 downregulated) comparing midlife low runners to midlife sedentary controls (**Fig. 3G)**. These results show that transcriptionally, running at a young age has a far greater effect on the number of DE genes than at midlife.

**Figure 3:**
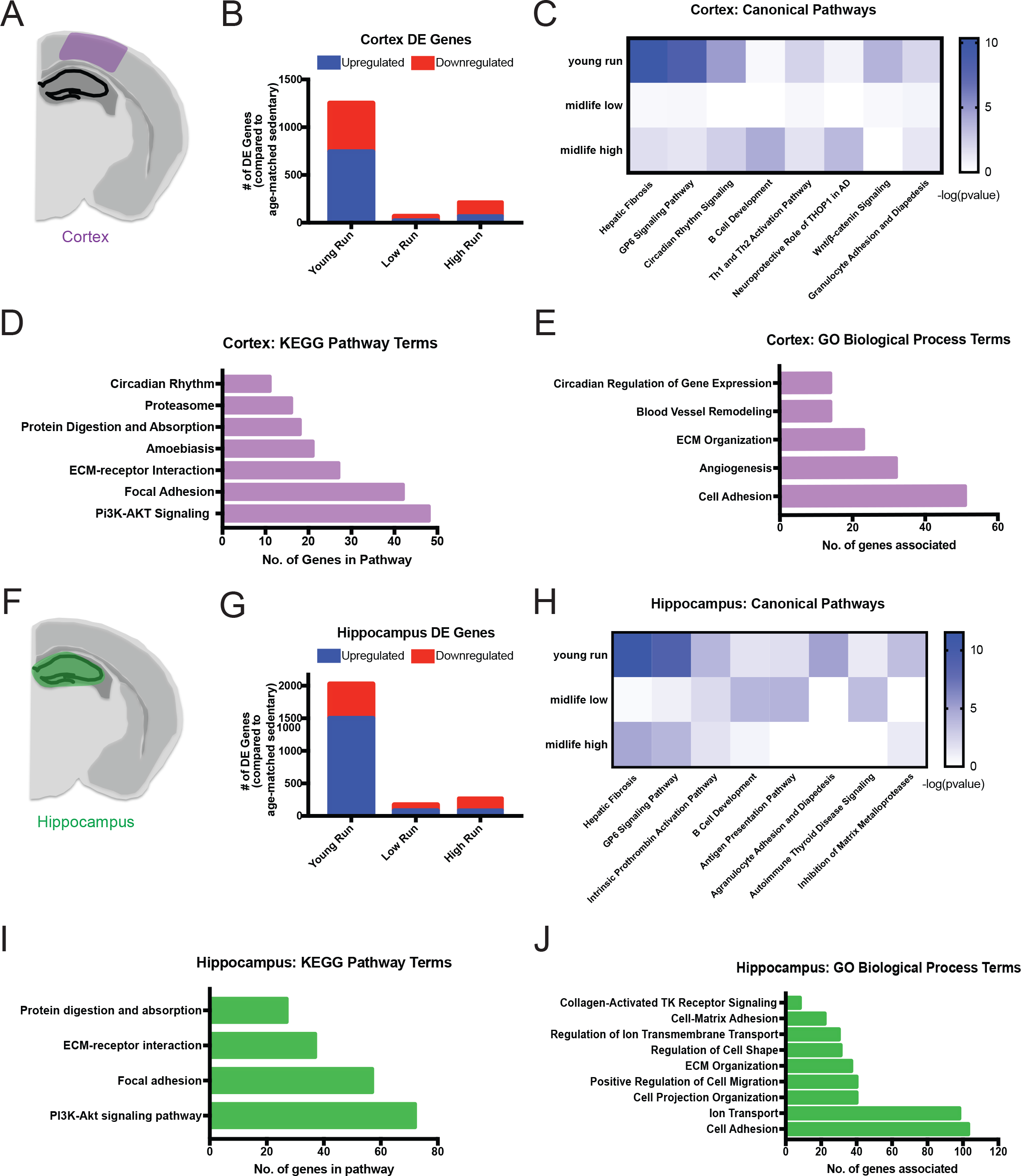
RNA-seq analysis identified ECM-related enrichment in both the cortex and hippocampus of young runners. (A) Cortical region (purple) used for RNA-seq. (B) Number of DE genes (FDR <0.05) found in the cortex of the young runners compared to young sedentary (‘Young Run’), midlife low runners compared to midlife sedentary (‘Low Run’), and midlife high runners compared to midlife sedentary (‘High Run’) (FDR<0.05). (C) IPA canonical pathway analysis showed enriched ‘Hepatic Fibrosis’ and ‘GP6 Signaling Pathway’ in the cortex of young run compared to young sedentary, however not significant in midlife comparisons. (D) KEGG pathway enrichment analysis in the cortex for young run compared to young sedentary (FDR<0.05). (E) GO term enrichment analysis in the cortex for young run compared to young sedentary (FDR<0.05). (F) Hippocampal region used for RNA-seq tissue submission. (G) Number of DE genes (FDR <0.05) found in the hippocampus of the young runners compared to young sedentary (‘Young Run’), midlife low runners compared to midlife sedentary (‘Low Run’), and midlife high runners compared to midlife sedentary (‘High Run’) (FDR<0.05). (H) IPA canonical pathway analysis showed enriched ‘Hepatic Fibrosis’ and ‘GP6 Signaling Pathway’ in the hippocampus of young run compared to young sedentary, however not significant in midlife comparisons. (I) KEGG pathway enrichment analysis in the hippocampus for young run compared to young sedentary (FDR<0.05). (J) GO term enrichment analysis in the hippocampus for young run compared to young sedentary (FDR<0.05).

### Extracellular matrix-related genes are altered in young but not midlife runners

Next, we sought to identify which pathways and genes were significantly altered by running at a young age, but showed no difference at midlife. We first analyzed transcriptional data through Ingenuity Pathway Analysis (IPA) canonical pathway analysis. DE genes comparing young runners to young sedentary controls were enriched for canonical pathway terms ‘Hepatic Fibrosis’, ‘GP6 Signaling’, and ‘Circadian Rhythm Signaling’ in the cortex and hippocampus. (**Fig. 3C, 3H)**. These pathways were not enriched in the midlife data. ‘Hepatic Fibrosis’ and ‘GP6 Signaling Pathway’ terms contain extracellular matrix (ECM)-related genes, such as collagens and laminins. KEGG pathway analysis further identified ‘ECM-receptor Interaction’ and ‘Focal Adhesion’ in the cortex and hippocampus, which substantiated the significant changes to the ECM (**Fig. 3D, 3I)**. There were no KEGG terms enriched in either the cortex or hippocampus for the midlife low runners. Only the KEGG pathway ‘Malaria’ was enriched in the cortex of midlife high runners. Genes DE in the ‘Malaria’ pathway are primarily hemoglobin-related genes suggesting a potential change in oxygenation in high runners. Only the KEGG pathway ‘ECM-receptor Interaction’ was enriched in the hippocampus of midlife high runners. Of the ten DE genes in ‘ECM-receptor Interaction’ were seven collagen or laminin genes (*Col24a1* (FDR= 2.39e-6, FC= −2.17), *Col6a3* (FDR= 0.026,FC= −1.38), *Col6a2* (FDR= 0.0032,FC=-1.45), *Col6a5* (FDR= 0.00029,FC= −1.72), *Col6a1* (FDR= 0.00025,FC= −1.52), *Col5a1* (FDR= 9.03e-8,FC= −1.75), and *Lamc2* (FDR= 6.85e-8,FC= −2.07)) which were all downregulated. This suggests that more intense running at midlife can influence ECM-related genes, but potentially not in a positive way.

GO terms are a way of categorizing DE genes into functional biological groups. GO term analysis of the DE genes from the cortex and hippocampus data from young runners showed enrichment for ‘ECM Organization’ and ‘Cell Adhesion’ **(Fig. 3E, 3J)**. Interestingly, vascular remodeling-related terms were significantly enriched in the cortex of young runners, including terms such as ‘blood vessel remodeling’, ‘ECM Organization’, ‘Angiogenesis’, and ‘Cell Adhesion’ **(Fig. 3E)**. None of these terms were enriched in the midlife data, irrespective of distance ran. However, one GO term, ‘Cellular Oxidant Detoxification’, which comprised mainly hemoglobin component genes, was enriched in both low and high runners in the hippocampus but only in the high runners of the cortex at midlife. These data suggest there is an overall change in ECM composition or organization due to running at a young age that is not reflected to the same degree at midlife. (**Fig. 3)**.

### Running upregulates genes related to vascular remodeling in young but not midlife mice

IPA revealed ‘Hepatic Fibrosis’ and ‘GP6 Signaling’ as significantly enriched canonical pathways in both the cortex and the hippocampus **(Fig. 3)**. These pathways contain many collagens and laminins that comprise the basement membrane component of the blood brain barrier that is important for maintaining brain health **(Fig. 4A)**. The majority of genes in ‘GP6 Signaling’ pathway were upregulated in the young runners in both the cortex and hippocampus but were not DE in either midlife high or low runners **(Fig. 4B-F)**.

**Figure 4:**
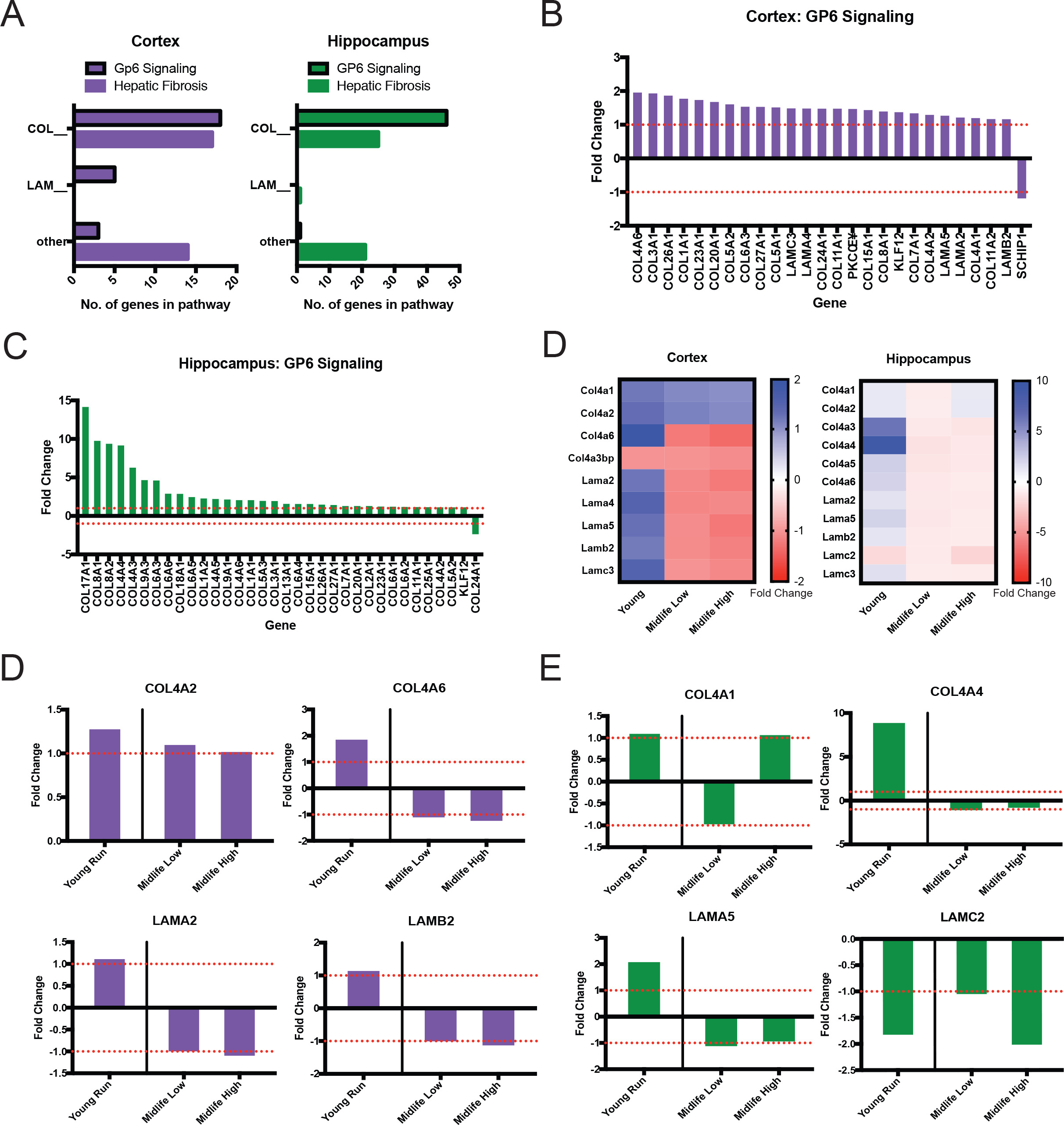
General upregulation of Collagens and Laminins due to young running with an attenuated response at midlife. (A) Number of significant collagen, laminin or other genes in young ‘Hepatic Fibrosis’ and ‘GP6 Signaling’ pathways in the cortex and hippocampus. (B) Fold changes of genes enriching for ‘GP6 Signaling’ in the cortex. (C) Fold changes of genes enriching for ‘GP6 Signaling’ in the hippocampus. (D) Heatmap of Col4s and laminins in the cortex and hippocampus that are significant in young run compared to young sedentary, while not significant in midlife comparisons. (E) Examples of significant cortical genes found in D showing attenuated responses in midlife cohort contrasts compared to young. (F) Examples of significant hippocampal genes found in D showing various responses in midlife cohort contrasts compared to young.

GO term analysis identified enrichment of ‘ECM organization’, ‘Blood Vessel Remodeling’ and ‘Angiogenesis’ in the cortex (**Fig. 5A**). These terms were significantly enriched in young mice but not in midlife mice. Of these vascular terms, ‘Angiogenesis’ contained 32 genes (26 upregulated, 6 downregulated) (**Fig. 5A**, B). At least six of the upregulated DE genes are implicated in *Vegf*-induced angiogenesis (*Mmp14* (FDR=1.23e-8, FC=1.75), *Vegfa* (FDR=0.038, FC=1.20), *Kdr* (FDR=.0011, FC=1.41), *Flt1* (FDR=.031, FC=1.21), *Dll4* (FDR=0.037, FC=1.59), *Notch1* (FDR= 0.002, FC=1.29)) (**Fig. 5C, 5D)** (19). However, of the 26 DE genes that were significantly upregulated in young runners compared to young sedentary, only two genes (*Col8a1*, *Hif3a*) were significantly DE in midlife high runners when compared to midlife sedentary controls (**Fig. 5C-D)**.

**Figure 5:**
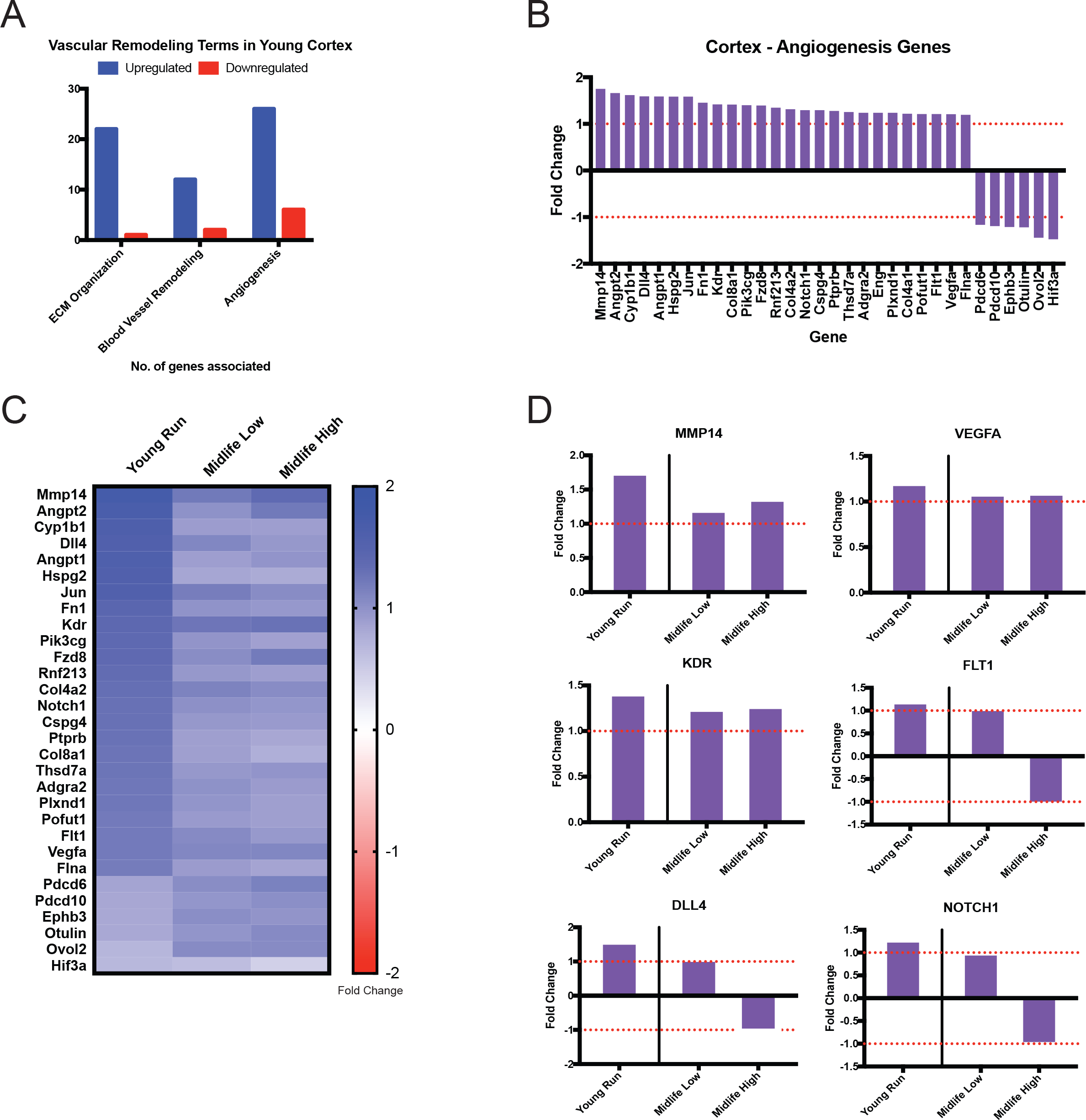
Angiogenesis genes are significantly enriched in young runners but not midlife runners. (A) GO terms enriched for vascular remodeling in the cortex show general upregulation. (B) Fold changes for DE genes enriched in the Angiogenesis term in the cortex. (C) Heatmap comparison between angiogenesis gene fold changes in the cortex in the young contrast (all significant) and midlife contrasts. (D) Examples of significant cortical angiogenesis genes found in C showing various responses in midlife cohort contrasts compared to young.

### Running activates genes in multiple cerebrovascular-related cell types

Finally, we determined whether specific cell types were more dramatically affected by young but not midlife running. This might provide insight into which cell type(s) are no longer responding to running in midlife mice. A sample of basement membrane and angiogenesis genes DE in young, but not midlife, running datasets were cross referenced to two cell-type specific datasets – the Brain RNA seq and a single cell RNA-seq dataset focused on cerebrovascular associated cells (20) (21). First, we evaluated *Col4* and laminin genes that were DE in young runners. As expected, the majority of the genes were expressed by cerebrovascular-related cells such as astrocytes and endothelial cells and not other cells in brain such as neurons, oligodendrocytes and microglia **(Table S9)**. However, many genes were expressed by different subsets of vascular-related cells such as *Col4a1* (endothelial cells, pericytes, vascular smooth muscle cells and fibroblasts) *Lamb2* (astrocytes, endothelial cells, pericytes, vascular smooth muscle cells and fibroblasts) and *Lama5* (astrocytes, endothelial cells and vascular smooth muscle cells) (**Fig. 6**). Second, we assessed cell type specific expression of angiogenesis genes. Upstream components of the angiogenesis pathway were mainly expressed by astrocytes, including *Mmp14* and *Vegfa* **(Fig. 7A**, B). Downstream components of the angiogenesis pathway were primarily expressed by endothelial cells (*Kdr*, *Flt1*, *Dll4*, *Notch1*). Therefore, ECM- and angiogenesis-related genes effected by running are expressed in multiple cell types relevant to the cerebrovasculature.

**Figure 6:**
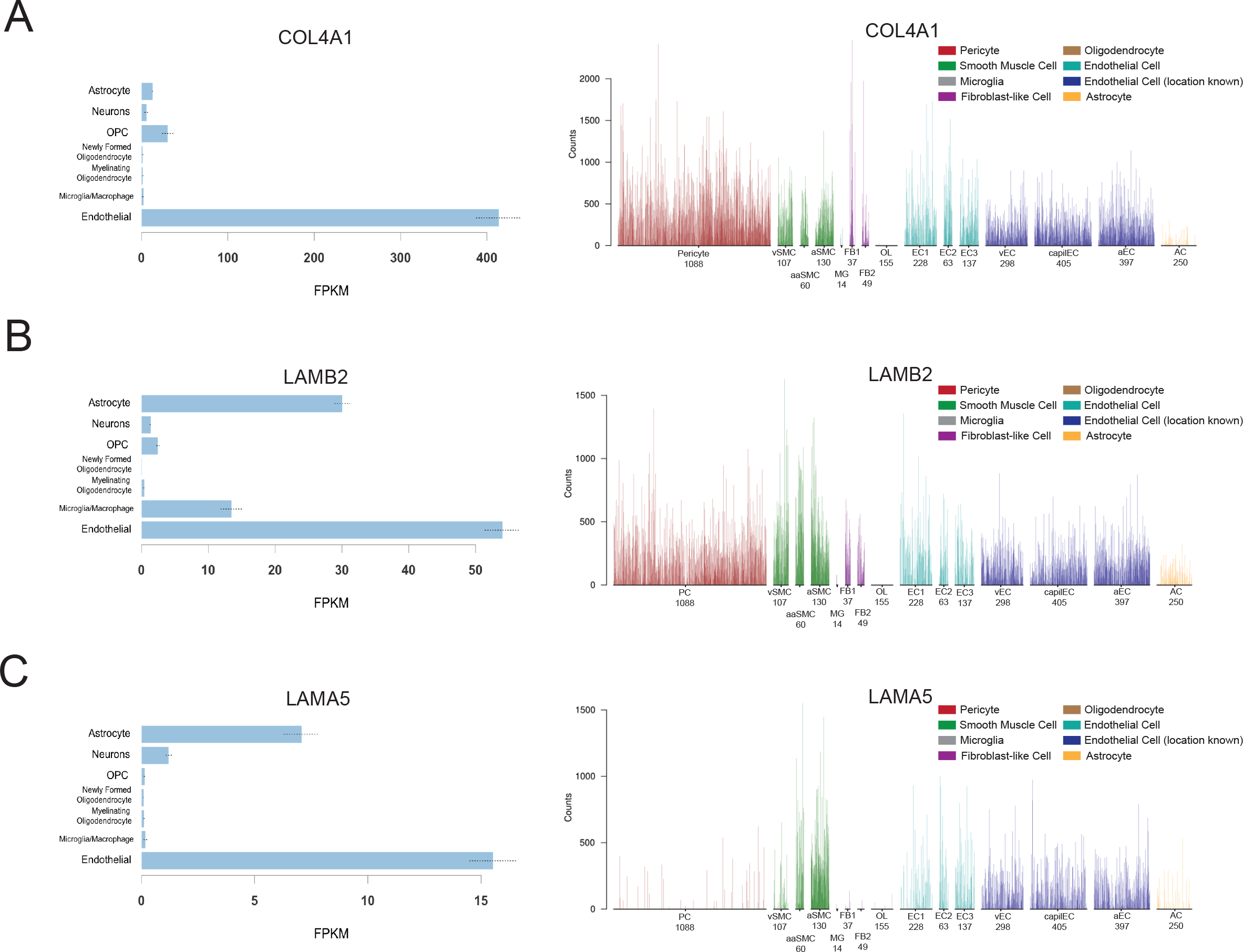
*Col4a1*, *Lamb2* and *Lama5* are expressed in multiple cerebrovascular-related cells. (A) Cell type specific expression from Zheng *et. al*. (left), and Vanlandewijck *et. al*. and He *et. al.* (right), showing overall expression of *Col4a1* in endothelial cells, smooth muscle cells, pericytes and fibroblast-like cells. (B) Expression of *Lamb2* in endothelial cells, smooth muscle cells, pericytes and astrocytes. (C) Expression of *Lama5* in endothelial cells and astrocytes.

**Figure 7:**
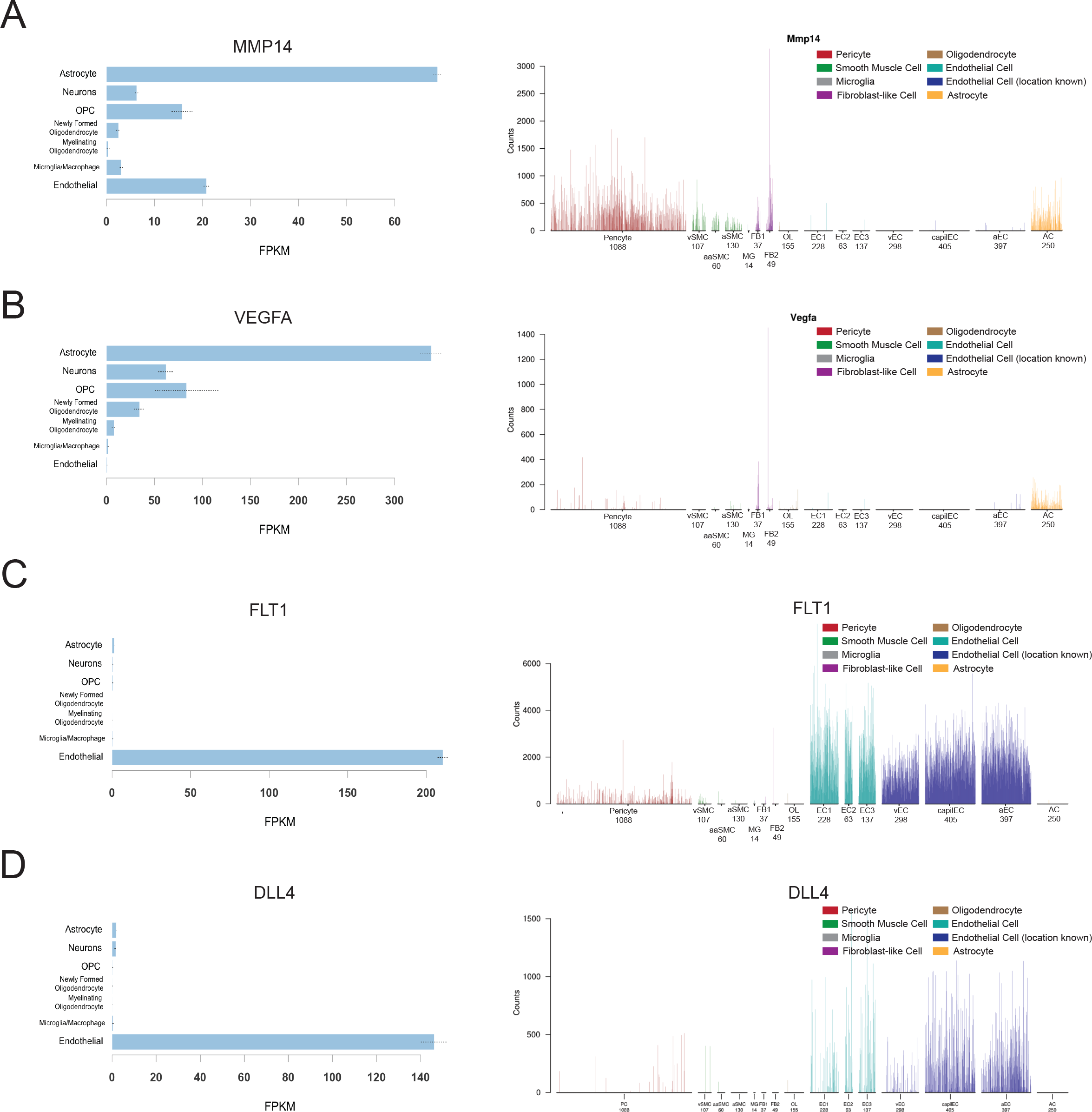
Angiogenesis genes are primarily expressed in astrocytes and endothelial cells. (A) Cell type specific expression from Zheng *et. al*. (left), and Vanlandewijck *et. al*. and He *et. al.* (right), showing expression of *Mmp14* in pericytes and astrocytes. (B) Expression of *Flt1* is primarily produced in endothelial cells. (C) Expression of *Dll4* is primarily produced in endothelial cells.

**Figure 8:**
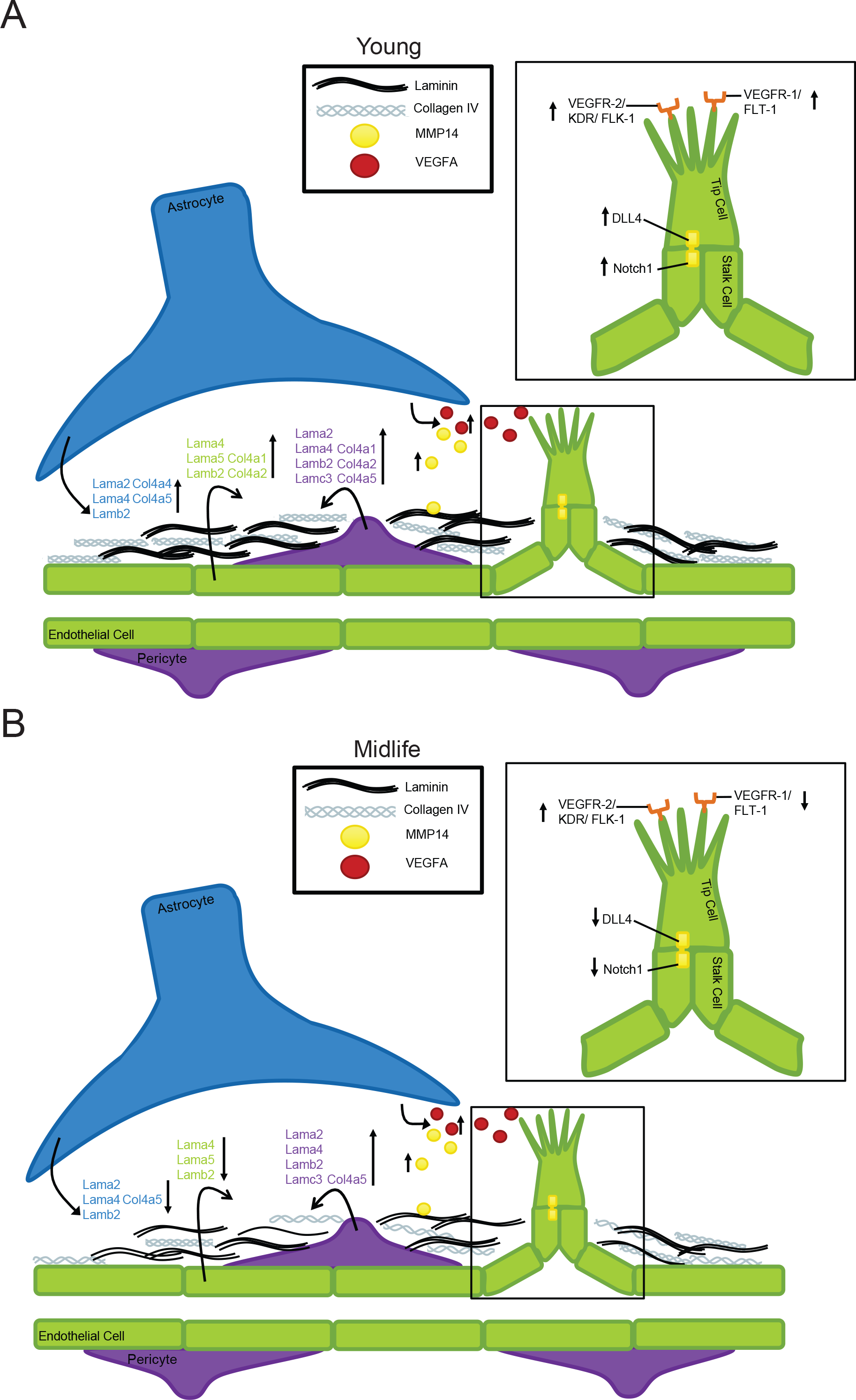
Running young is more effective than running at midlife. (A) Depiction of transcriptional changes occurring in a young mouse due to running, including upregulation of collagens and laminins from astrocytes, pericytes and endothelial cells, as well as induction of the VEGFA-KDR-DLL4 angiogenesis pathway. (B) Depiction of molecular changes occurring in midlife mice due to running, including downregulation of collagens and laminins, and a trend for induction of early angiogenesis genes from astrocytes (MMP14, VEGFA) but an attenuated downstream angiogenesis response (FLT-1, DLL4, NOTCH1) in endothelial cells.

## Discussion

Here, we assessed B6 male mice at two ages, young and midlife, to better understand the effects of running on the aging brain. Our data suggest that intervention of running may not be as beneficial to the cerebrovasculature in midlife. While systemic health (lipid profile, body composition) is improved by running at midlife, the cerebrovasculature is not as responsive. Therapies or interventions during midlife may require different approaches than preventative measures in young, more responsive individuals. In this study, half the midlife cohort (high runners) ran as much as the young cohort, whereas the other half ran far less (low runners, **Fig. 1**). Interestingly, this did not drastically affect the transcriptional profiles (**Fig. 3**). We would have anticipated that running further and faster would have been more beneficial at midlife, however, our results show a mitigated response in all midlife runners compared to young runners.

Previous exercise studies in mouse models have primarily focused on the benefits of exercise in young rodents. Findings from these experiments are promising, showing that running can reduce amyloid plaque development, induce neurogenesis, reduce infarct volume, and improve cognition (14, 22-24). However, the brain during development and early adult life is considered more plastic than in later life, which incites therapies that will not properly translate to aged trials (2) (25).

We took an unbiased approach to identify transcriptional changes in two vulnerable brain regions (cortex and hippocampus) as a result of running. We show that the greatest change from running was to the cerebrovasculature. Particularly, we highlight genes involved in basement membrane composition (particularly collagens and laminins) that are upregulated in young mice due to running that were not affected by running at midlife. Similarly, genes in the angiogenesis pathway (e.g. *Mmp14*, *Vegfa*, *Kdr*, *Flt1, Dll4,* and *Notch1*) showed induction by running in young but not midlife mice (19). These transcriptional data suggest that the benefits of exercise to the cerebrovasculature declines with age, increasing the need for a greater understanding of midlife exercise as an intervention for cognitive decline and dementia (26).

Previous studies have focused on the neuroprotective benefits of exercise. In Choi et al., of the 5XFAD mice (a model relevant to AD) that were exercised, half were categorized as showing increased neurogenesis while the other half did not show this effect, potentially due to the amount run (14). Additionally, it was shown that viral-induced neurogenesis was not enough to rescue cognitive decline, while exercise-induced neurogenesis was able to sufficiently improve cognitive results (14). This finding suggests that running is providing additional benefits to substantiate neurogenesis mediated improvements in cognition. Based on the results of our study, we predict that in Choi *et al.* there were running-induced improvements to the cerebrovasculature which would provide better clearance of amyloid and better maintenance of neuronal health resulting in improved cognition (14). It is known that the development of the neuronal and vasculature systems is strongly linked, with growth occurring simultaneously because neurons require vascular support for oxygen and nutrients. This further supports the need for cerebrovascular improvement to accompany adult therapy-induced neurogenesis (27). If neurogenesis and angiogenesis become uncoupled, this may cause stress to neurons, reduced cerebral blood flow, and a disruption to neurovascular coupling. Therefore, when considering midlife interventions to cognitive decline and dementia it is important to consider both neuronal and cerebrovascular health.

A limitation of our study is that we used healthy wild-type male mice to assess the effects of exercise at two different ages. In our study, the effects of exercise were greater in young compared to midlife male mice. It is possible that exercise at midlife would be beneficial in female mice and also in mice that are predisposed to vascular damage and dementia through genetic or environmental risk factors. Due to the increasing recognition of a vascular contribution to dementia symptomology, it would be pertinent to know if running alleviates these effects of vascular risk in genetically predisposed mice at different ages. For example, the ε*4* allele of apolipoprotein E *(APOE*^*ε4*^), the greatest risk factor for late-onset AD and Vascular Dementia, shows early cerebrovascular decline in both mice and humans (28) (29). This is proposed to be due to a lack of binding of *APOE*^*ε4*^ to the low density lipoprotein receptor related protein 1 (*LRP1*) leading to an increase in *MMP9* and a breakdown of basement membrane proteins and endothelial cell tight junctions (28). Therefore, when evaluating running as a potential intervention, it is imperative to understand whether benefits will be seen across multiple dementia risk genotypes. Although these experiments are challenging to perform in human populations, they can be readily performed in mouse models.

Due to the heterogeneity of dementia pathology, it is possible that the typical approach to treating dementias, is too narrow in scope, only able to alleviate the burden of a small subset of dementia cases. Only recently has the American Heart Association acknowledged the prevalence of Vascular Contributions to Cognitive Impairment and Dementia (VCID), explaining that cerebral infarcts are frequent and common with age, including in patients with diagnosed with AD (30). We hypothesize that exercise is key in identifying new pathways, specifically those related to cerebrovascular health, that may help a broader population of dementia patients. In summary, our research and others show that exercise benefits cerebral health by improving multiple systems including the cerebrovasculature. However, the benefits of exercise appear to decline with age, further supporting that combinatorial approaches are required to prevent cognitive decline and dementias.

## Abbreviations

AD: Alzheimer’s Disease
VCID: Vascular Contributions to Cognitive Impairment and Dementia
B6: C57BL/6J
mo: months
BDNF: Brain derived neurotrophic factor
DE: Differentially expressed
DGE: Differential gene expression
HDL: High density lipoprotein
LDL: Low density lipoprotein
NEFA: Non-essential fatty acids
ECM: Extracellular matrix
Hippo: hippocampus
Ctx: cortex
NMR: nuclear magnetic resonance
RNA-seq: RNA sequencing
FDR: False Discovery Rate
KEGG: Kyoto Encyclopedia of Genes and Genomes
DAVID: Database for Annotation, Visualization, and Integrated Discovery
GO: Gene Ontology
IPA: Ingenuity Pathway Analysis
ANOVA: Analysis of Variance

## Declarations

### Author’s Contributions

KEF, LCG and GRH conceived and designed the project. KEF and LCG performed experiments. KEF and SY performed bioinformatic analysis. KEF and GRH wrote this manuscript. All authors approved this manuscript.

## Acknowledgments

The authors wish to thank Todd Hoffert from Clinical Assessment Services for blood chemistry, Heidi Munger and the Genome Technologies group for RNA-sequencing, and Tim Stearns and Vivek Philip from Computational Sciences.

## Availability of Data and Resources

Upon acceptance all data will be available on GEO archive.

## Funding

This work was supported by T32HD007065 (KEF) and by a kind donations from Tucker and Phyllis Taft (GRH).

## Competing interests

The authors declare they have no competing interests.

## Ethics approval and consent to participate

Not applicable

## Consent for publication

Not applicable

