## Supplementary material for "Transcriptional profiling reveals running promotes cerebrovascular remodeling in young but not aged mice": Supplmental Figures and Legends

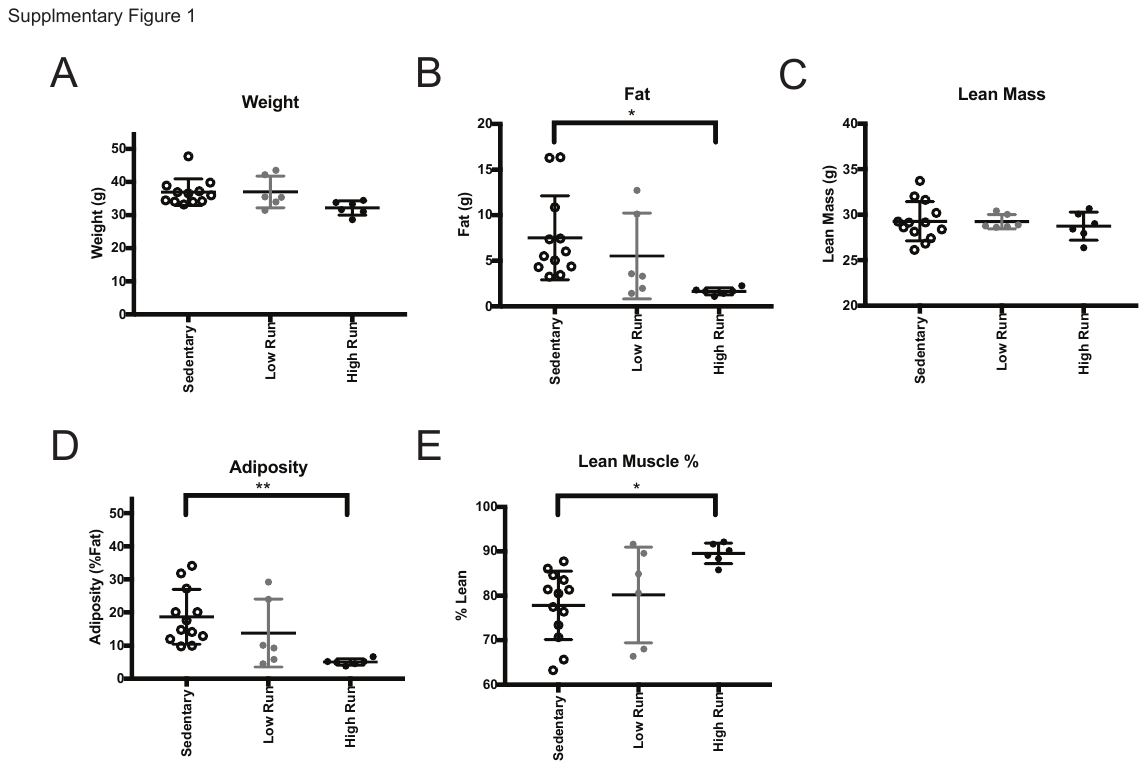

**Supplemental Figure 1: Midlife High runners show reduced adiposity due to running.** (A) Weight of midlife mice at time of NMR showed no significant difference. (B) Significant reduction in amount of fat (grams) in midlife high compared to midlife sedentary mice (*p =0.0225). (C) No significant difference in lean mass (grams) between the midlife cohorts. (D) Significant reduction in adiposity (%fat) in midlife high runners compared to midlife sedentary (**p= 0.0060). (E) Significant increase in lean muscle percentage of midlife high runners compared to midlife sedentary (*p= 0.0154).

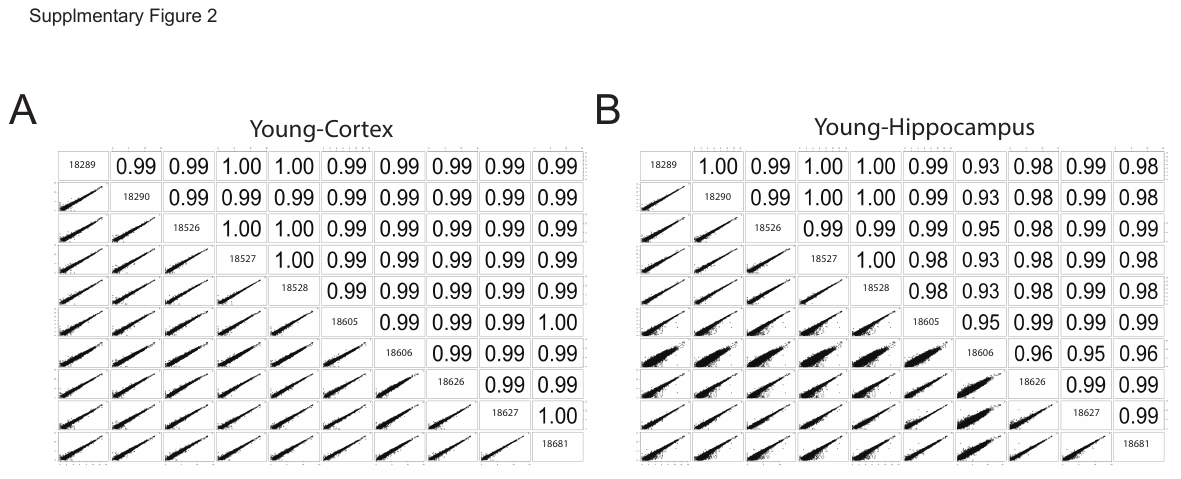

**Supplemental Figure 2: CPM scatterplot of young runners compared to young sedentary in the cortex and hippocampus.** (A-B) Counts per million (CPM) scatterplot of young runners compared to young sedentary in the cortex (A) and hippocampus (B) revealed Mouse 18606 as an outlier from the hippocampal dataset. This sample was removed from further analysis.

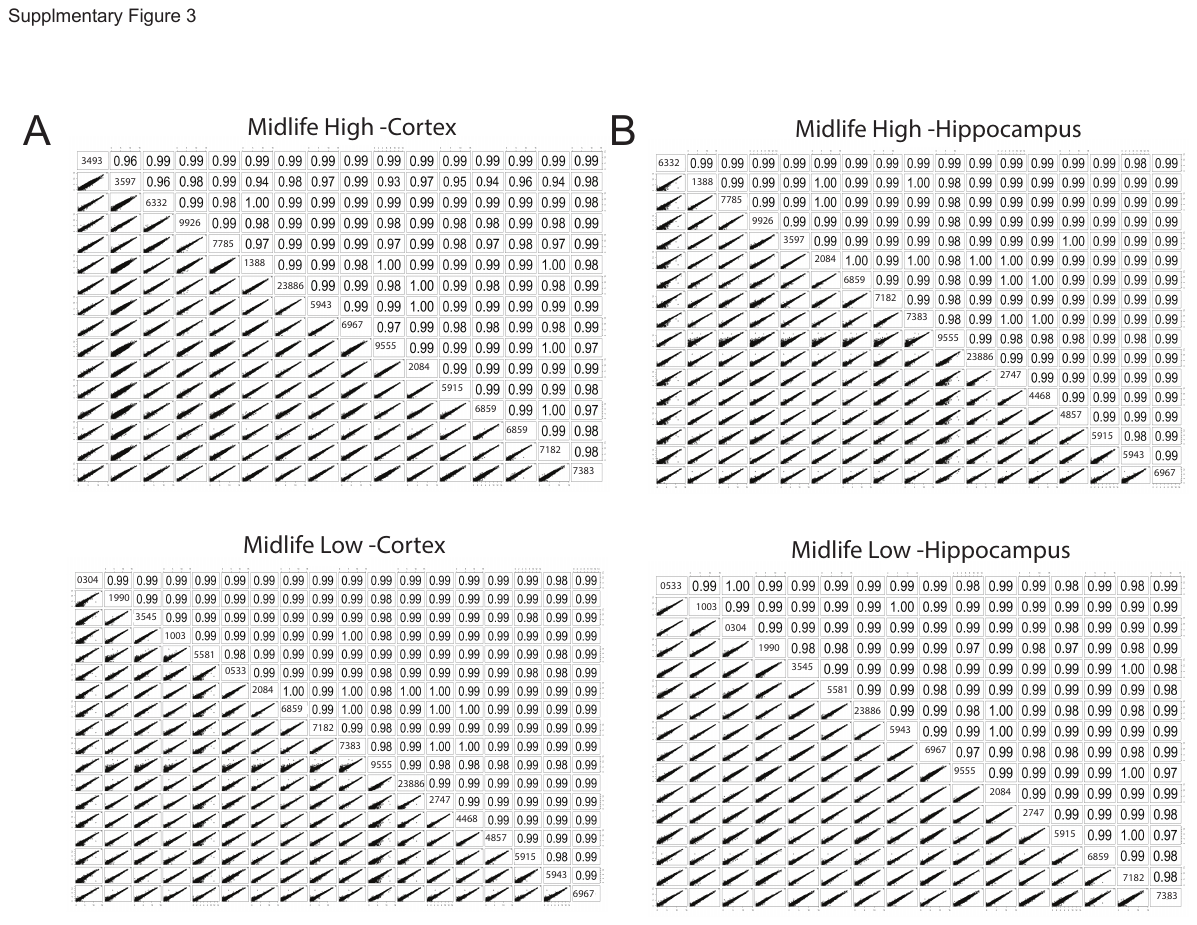

**Supplemental Figure 3: CPM scatterplot of midlife runners compared to midlife sedentary in the cortex and hippocampus.** (A-B) Counts per million (CPM) scatterplot of midlife runners compared to midlife sedentary in the cortex (A) and hippocampus (B) revealed no outliers in this dataset. There was also no observable difference between high runners and low runners.

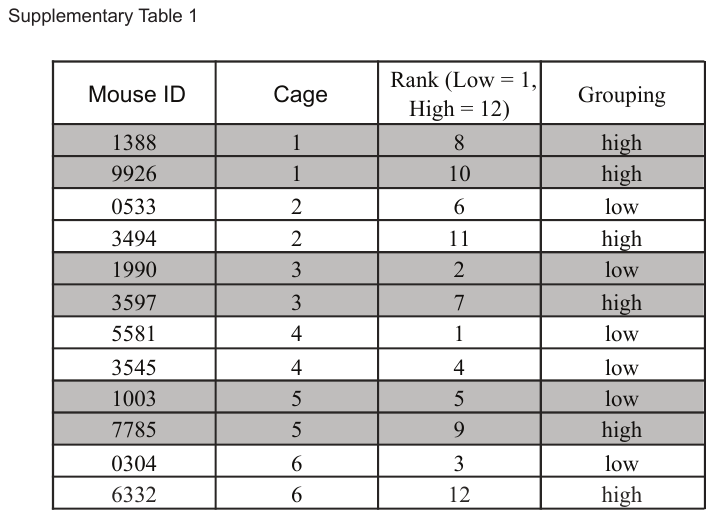

**Supplemental Table 1: Housing of Midlife Running mice.**

**Supplemental Table 2: Background Gene Sets for DAVID KEGG/GO term analysis**

See separate excel files.

**Supplemental Tables 3-8: DE gene lists from young and midlife high and low comparisons**

See separate excel files.

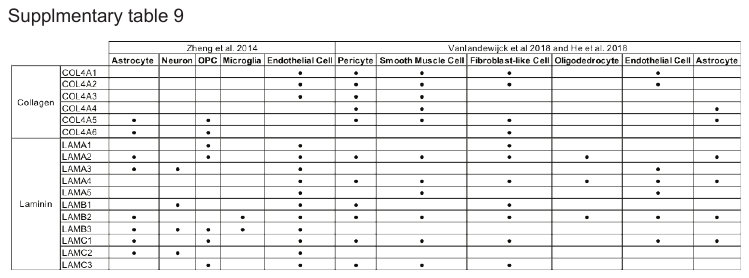

**Supplemental Table 9: Expression of Laminins and Collagen IV genes in both Zheng et al, and Vanlandewijck and He et al.**
